# C-Jun N-terminal kinase (JNK) pathway activation is essential for dental papilla cells polarization

**DOI:** 10.1101/2020.05.18.101782

**Authors:** Jiao Luo, Xiujun Tan, Ling Ye, Chenglin Wang

## Abstract

During tooth development, dental papilla cells could develop into odontoblasts with polarized morphology and cell function, c-Jun N-terminal kinase (JNK) signaling could participate in this process. Histological staining, qPCR and Western Blot shown that activation of JNK signaling in polarized mouse dental papilla tissue. *In vitro* cell culture and organ culture method found JNK inhibitor SP600125 postponed tooth germ development and reduced the polarization, migration and differentiation of mouse dental papilla cells (mDPCs) *in vitro*. The expression of polarity-related genes including Prickle3, Golga2, Golga5 and RhoA was consistent with JNK signaling activation, by screening of up-regulated polarity-related genes during the process of dental papilla development and mDPCs or A11 differentiation. Further, constitutively active RhoA mutant (RhoA Q63L) partly rescue the inhibition of SP600125 on cell differentiation and polarity formation of mDPCs. This study suggests that JNK signaling has a positive role in dental papilla cells polarization formation.

## Introduction

At early stages, dental papilla cells are small with centered nuclear and a few organelles, then appear to differentiate into preodontoblasts which located randomly along basement membrane (BM) and differentiate into polarizing odontoblasts which are tall columnar with polarized distribution of nuclear and cytoplasmic organelles[1, 2]. Study about dental papilla cells polarization is helpful to further understand the molecular mechanism in tooth development and pulp repair.

Few studies have tried to explore the molecular mechanism during the process of dental papilla cells polarization. For example, epithelial basolateral maker aquaporin 4 and apical aquaporin 5 were located at the plasma membrane of odontoblast processes or cell surface toward pulp in functional odontoblasts, respectively [3]. In transgenic mice, defective odontoblast polarization such as apical nuclei distribution was found in Epfn (-/-) mice[4], and dentin formation was disrupted in DLX3 mutant mice[5]. Several polarity proteins specifically localized in the odontoblastic layer such as MAP1B[6], PRICKLEs[7], Celsr1[8]. JNK signaling is activated primarily by cytokines such as tumor necrosis factor, epidermal growth factor, TGF, or exposure to diverse environmental stresses. By JNK-/- mice, C-Jun N-terminal kinase (JNK) kinase maybe participating in tissue morphology as spine [9]and in expression of four polarity complex genes as Celsr2, Numb, Prkcz, Llgl2 in bone marrow hematopoietic progenitor cells[10]. Previous studies showed JNK signaling was participated in cell proliferation, differentiation, apoptosis, and cytoskeleton reorganization[11–13], and regulating plane cell polarity(PCP)[14]. Our previous study[15] had found JNK pathway could regulate human dental papilla cells adhesion, migration, and formation of focal adhesion complexes, which was related with cell polarization, so here we further study the role of JNK pathway in dental papilla cells polarity formation.

## Materials and methods

### Tissue samples collection

All procedures performed in this study were in accordance with the ethical standards of the Ethical Committees of West China School of Stomatology, Sichuan University. ICR/CD1 mice were housed in small groups in specific pathogen free conditions at 22°C with a 12h light/ dark cycle and ad libitum access to food and water. The pregnant mice were culled by CO2 inhalation after successful mating for some days, then cardiac perfusion with saline performed before cervical dislocation and collection of tooth germ tissue. The heads or mandibles with tooth germs of ICR/CD1 mice were isolated, at embryonic day 17.5(E17.5), E18.5(vaginal plug = E0.5) and postnatal day 1 (P1), then fixed in 4 % paraformaldehyde (PFA), embedded in paraffin, and cut into 5μm sections. Hematoxylin and eosin(H&E) staining and immunofluorescence (IF) or immunohistochemistry (IHC) staining were performed.

### Immunofluorescence and immunohistochemistry

The slides were dewaxed in xylene, rehydrated and then subjected to antigen retrieval in citrate buffer at 99 °C for 5 min×3. For IF, the primary antibody anti-GM130 (Golgi matrix protein 130) (1:10, BD Pharmingen, USA) was incubated overnight at 4°C. Nuclear was stained using DAPI. For IHC, the primary antibodies, anti-phospho-JNK (Tyr183/Tyr185) or anti-JNK (1:50, Cell Signaling Technology Inc., MA, USA) were incubated overnight at 4°C, and the next procedures were performed with SP9001 kit and DAB Staining Kit (ZSGB-Bio, Beijing, China).

### Quantitative real-time PCR and western bolt analysis

Before RNA extraction and protein extraction, E17.5 and P1 tooth germ should be separated into mesenchymal and epithelial tissue, and the separation methods were performed as previously described[16].

Total RNAs were isolated from E17.5 and P1 separated tooth epithelial and mesenchymal tissues using TRIzol Reagent (Life Technologies, Carlsbad, CA, USA) according to manufacturer’s protocol. After cDNA was synthesized using the PrimeScript RT Reagent Kit (Takara, Dalian, China) and measured using a CFX96™ real-time PCR detection system (Bio-Rad, USA) as described previously[17]. Primer sequences are searched from PrimerBank (S1 Table). Relative gene expression levels are presented as the mean and standard deviation from three independent experiments.

Total proteins of E17.5 and P1 tooth germ mesenchymal tissues were isolated, and protein concentration was measured, protein transfer and incubate following regular methods. Anti-GAPDH, anti-phospho-JNK (Tyr183/Tyr185) (1:1000), anti-GM130 (1:500), and HRP-conjugated anti-rabbit secondary antibodies (1:1000, Cell Signaling Technology Inc., MA, USA) were used. Finally, we determined the ratio of the target protein to the reference protein. The obtained ratios are presented as the mean and standard deviation from three independent experiments.

### Tooth germ organ culture *in vitro*

First, mandibular first molar tooth germs from E16.5 CD1 mouse embryos were harvested under dissection microscope following conventional procedures[16, 18]. Second, the separated tooth germs were cultured in a modified Trowell-type organ culture in a 12-well 8.0 μm Transwell (BD Biosciences, USA) and grown at the air-liquid interface[19, 20], with or without 15μM SP600125 (Sigma, St. Louis, MO, USA) in cultured medium. Then, the culture plates were kept in a humidified incubator at 37°C in 5% CO2 for 5 or 7 days, with medium change every 2 days, before the tooth germs were rinsed and fixed in 4% ice cold-PFA, embedded in paraffin, and cut into 5μm sections. H&E staining was performed to examine tissue and cell morphology.

### Cell culture and JNK signaling inhibition

The mouse dental papilla cells (mDPCs) were cultured from separated dental papilla tissues of P1 tooth germs following digestion with 3 mg/ml collagenase I (Sigma, St. Louis, MO, USA) for approximately 45min at 37°C. The tissue piece and cell suspension were seeded into 60-mm cell culture dishes and cultured in Dulbecco’s modified Eagle’s medium (DMEM, Corning, USA) supplemented with 15% fetal bovine serum (FBS, Gibco, USA) and 1% penicillin/streptomycin (P/S, Hyclone, USA). Upon 90% confluence, the mDPCs were collected and sub-cultured for use. Mouse odontoblast-like A11 cells line was maintained in DMEM supplemented with 10% FBS and 1% P/S at 37°C and 5% CO2.

MDPCs and A11 cells were seeded at the same density and pre-incubated with 10μM SP600125 for 2h or 24h at 37°C before scratch assay or regular osteogenic medium (OM) treatment. Medium, inhibitor, and osteogenic medium were freshly added every second day.

### Scratch assay and cell immunofluorescence

The scratch assay and cell IF staining were performed as previously described[15, 21, 22]. Scratches were generated using a yellow plastic pipette tip after treated with 10μM SP600125 for 2h. Following treatment with SP600125 (10μM) within 24h, an inverted microscope (Diaphot, Nikon Corporation, Tokyo, Japan) was used to take wound healing images. Relative rates of cell migration were measured and expressed as a percentage of the initial length at zero time. Next, fixed cells were incubated with the primary antibodies, anti-GM130(1:10), and the secondary antibody, anti-rabbit IgG FITC conjugated (1:200, Santa Cruz, CA, USA). Cells at the leading edge of scratch line were observed, with anti-GM130 labeled Golgi apparatus distribution. Quantitation of cells at leading edge with Golgi polarized within a 120°arc in front of nucleus was analyzed. Each experiment was repeated three times.

### Alkaline phosphatase (ALPase) and alizarin red S (ARS) staining

As the change of cell functional polarity was hard to direct detect *in vitro*, and cell functional polarization will lead to cell differentiation, so we had to use the change of cell differentiation ability to present the change of cell functional polarity. In this study, we use ALPase staining and ARS staining to check the change of cell differentiation of mDPCs and A11 cells. For ALPase staining, cells were cultured in OM with or without 10μM SP600125 for 3 or 5 days and tested by ALPase staining kit (Beyotime Institute of Biotechnology, Hai-men, China) according to manufacturer’s instructions. For ARS staining, cells were cultured in OM with or without 10μM SP600125 for 7 or 14 days and tested by ARS staining (Sigma-Aldrich) according to manufacturer’s instructions. Each experiment was repeated 3 times.

### Polarity-related gene screening

Total RNAs were isolated from E17.5 and P1 dental papilla tissue, from mDPCs before and 6h after OM induction, from A11 cells before and 6h after OM induction. Then cDNA was synthesized and real-time PCR was performed as described above. Primers of some polarity-related genes were also in S1 Table.

After we screen out some up-regulated polarity-related genes during the process of dental papilla development and mDPCs or A11 differentiation. The expression of these screened genes was confirmed in SAPK/JNK siRNA transfected or SP600125 treated mDPCs. For siRNA transfection, the cells were transfected with 100nM SAPK/JNK siRNA I or control siRNA (Cell Signaling Technology Inc., MA, USA) by using Lipofectamine 3000 (Invitrogen, life technologies, USA) according to the manufacturer’s instructions. For SP600125 treatment, the cells were incubated with 10μM SP600125 for 24h at 37°C. Then RNAs were isolated and qPCR was performed as above.

### RhoA mutant adenovirus construction and transfection

Our previous study[15] had constructed RhoA mutant recombinant adenovirus including Ad-RhoA WT (wild type RhoA) and Ad-RhoA Q63L (constitutively active RhoA mutant), and transfected them into human dental papilla cells. This study [15] suggested both RhoA and JNK signaling had roles in regulating cell adhesion and migration. So here, RhoA was further chose to study its role in JNK signaling regulated cell polarity. First, mDPCs were transfected by Ad-RhoA WT or Ad-RhoA Q63L for 48h. Before ALPase staining as above, mDPCs were cultured in OM with or without 10μM SP600125 for 5 days. Before scratch assay and IF staining as above, mDPCs were culture in regular medium with or without 10μM SP600125 for 24h. Each experiment was repeated 3 times.

### RhoA pull-down assay

Ad-RhoA WT or Ad-RhoA Q63L were transfected into mDPCs for 48 h, following SP600125 treatment for 24 h, then GST Pull-down assay with a glutathione transferase (GST) fusion protein containing the RhoA binding domain of rhotekin (rhotekin-GST) was performed by the manufacturer’s protocol using GTPase Pull-Down kit (Thermo Scientific, Waltham, MA, USA). Activated and total RhoA were detected by Western blot analysis using anti-RhoA antibody[15].

### Statistical analysis

All values were calculated as the mean ±standard deviation (SD). The statistical significance was determined by unpaired Student’s t-test of variance at *P*<0.05.

## Results

### Dental papilla cells polarization during tooth germ development

Dental papilla cells, especially those close to BM, gradually underwent various development stages, the cells were defined as proliferation zone, differentiation zone and secretory zone from apical to coronal[7, 23]. At incisor, dental papilla cells were small and undifferentiated in proliferation zone (Fig 1, A1), columnar with polarized distribution of nuclear in differentiation zone (Fig 1, A2), functional odontoblasts with matrix synthesis and secretion function in secretory zone (Fig 1, A3). The same phenotype was repeated at cusps region of first molar (Fig 1, B-D), dental papilla cells show different morphology at different stage, similar to incisor. Meanwhile, IF staining showed Golgi marker GM130 were also gradually orderly distributed at the function side close to BM in differentiation and secretory zone, also suggesting cell polarity formation in these zones (Fig 1, E). In detail, GM130 were laterally distributed in the functional side of odontoblasts (Fig 1, E2, E2’, E2’’), but not cervical loop region (Fig 1, E1, E1’, E1’’).

**Fig 1.**
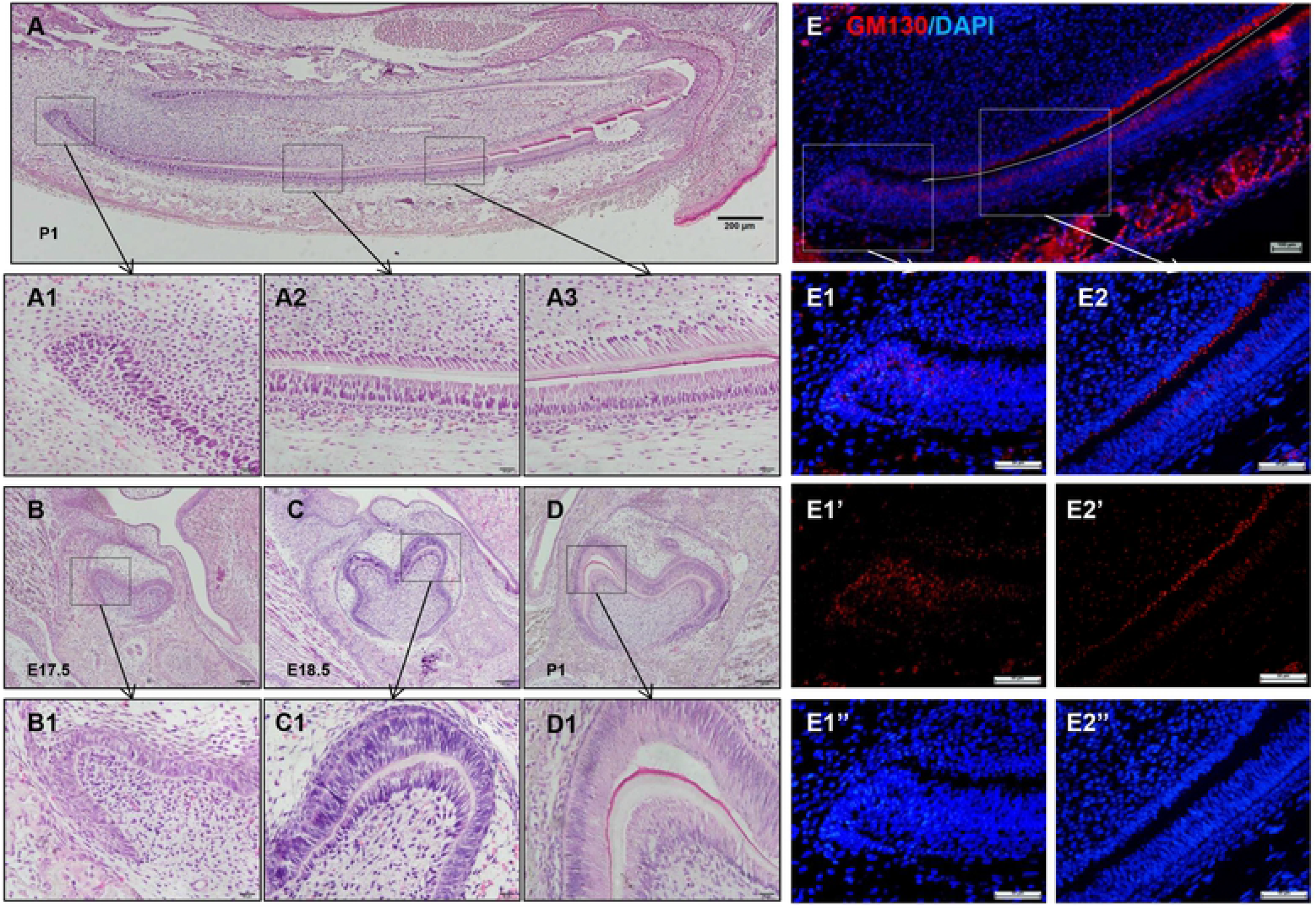
Morphogenesis and function of dental papilla cells polarization. (A) H&E staining of P1 incisor. From apical to coronal: A1, Proliferation zone. Small and undifferentiated. A2, Differentiation zone. Columnar and cell polarity morphogenesis. A3, Secretory zone. Odontoblasts-like with cell polarity function. (B-D) H&E staining of the mandibular first molars from E17.5, E18.5 and P1. B1-3, At cusps region, dental papilla cells changed from flat shape to columnar with polarity formation, and high columnar. (E) IF of GM130 at P1 incisor. GM130 were laterally distributed in the functional side of odontoblasts (E2, E2’, E2’’), but not cervical loop region (E1, E1’, E1’’). E: embryonic, P: postnatal, Bars: 200μm (A), 20μm (A1-A3, B1-D1), 100μm (E), 50μm(E1-E2’’)

### Activation of JNK signaling in polarized dental papilla cells

In P1 incisor, p-JNK was mainly expressed in differentiated and secretory odontoblasts (Fig 2, A1-A3), while total JNK was widely expressed in odontogenic cells (Fig 2, B1-B3). In molar, p-JNK was barely visible in dental papilla at E17.5 (Fig 2, C), but strongly expressed in odontoblasts at P1(Fig 1, D). The results showed that the spatial and temporal expression of JNK signal had a correlation with dental papilla cells polarization.

Under the stereomicroscope, we separated the dental epithelium and dental papilla from the mandibular first molars of E17.5 and P1(S1 Fig A, B), and detected the expression of epithelium marker Cdh1 and mesenchymal marker Lhx8, which simply verified separated tissue source properties and availability (S1 Fig C). RT-PCR showed increased expression of Jnk1 and Jnk2 in P1 dental papilla compared with E17.5, also GM130/Golga2 (Fig 2, E). Western blot showed increased expression of p-JNK and GM130 in P1 dental papilla compared with E17.5 (Fig 2, F). These results suggested JNK signaling was activated when dental papilla cell polarity formed.

**Fig 2.**
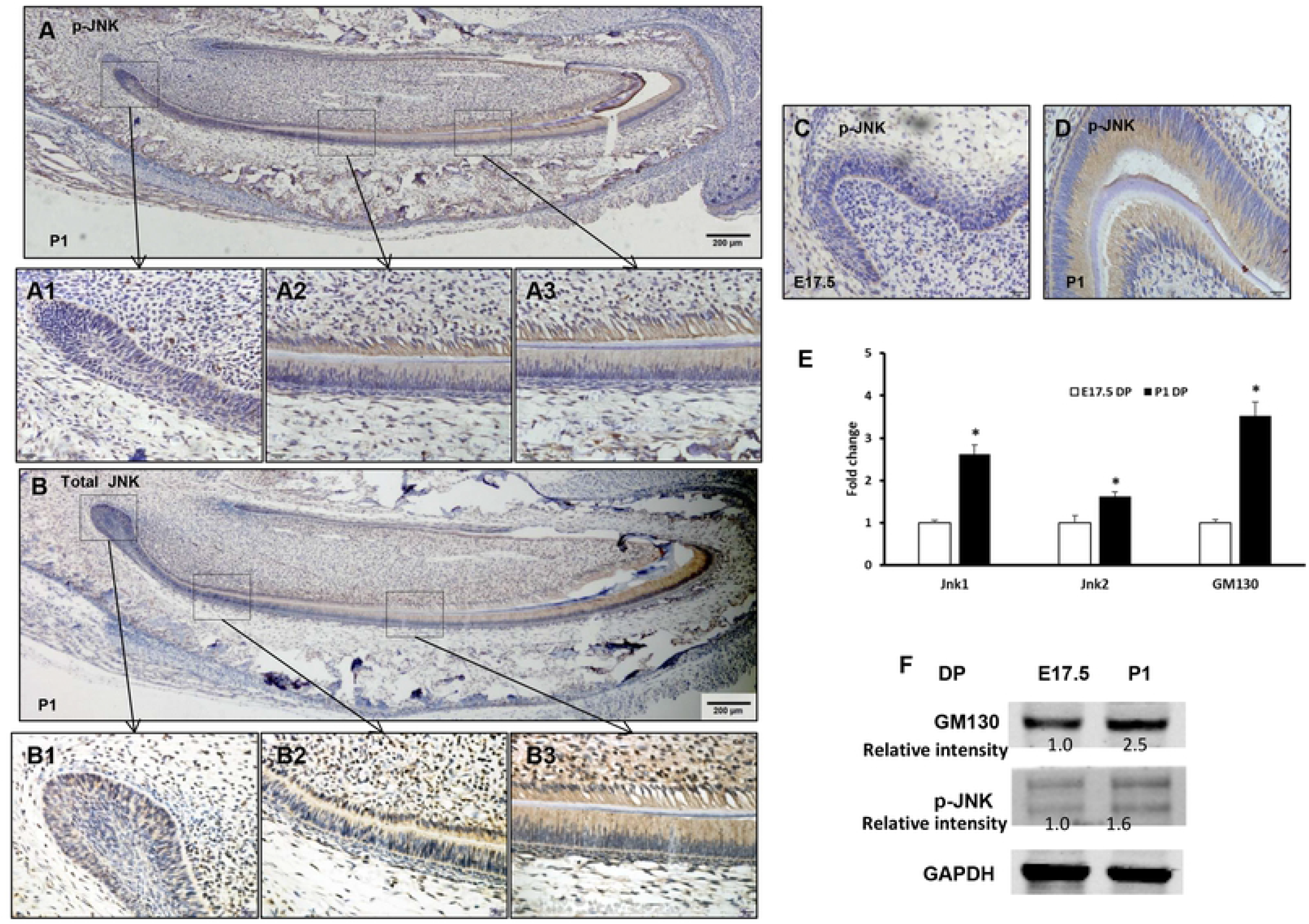
Activation of JNK signaling in polarized dental papilla cells. (A) At P1 incisor, immunochemistry revealed that p-JNK was mainly expressed in differentiation zone(A2) and secretory zone(A3), but not in proliferation zone (A1). (B) Total JNK was widely expressed in odontogenic cells(B1, B2, B3). (C) At E17.5, p-JNK was barely visible in dental papilla. (D) At P1, p-JNK was strongly expressed in odontoblasts. (E) RT-PCR showed increased expression of Jnk1, Jnk2 and GM130 in P1 dental papilla tissue, compared with E17.5. (F) Western blot analyses of p-JNK and GM130 in E17.5 and P1 dental papilla tissues. DP: dental papilla, Bars: 200 μm (A,B), 20 μm (A1-A3,B1-B3,C,D)

### The effect of JNK inhibitor on tooth germ development *in vitro*

Given the JNK signaling was playing a role in tooth germ development, we wondered whether JNK inhibitor SP600125 could induce retardation of tooth development. Concentration screening showed 10μM and 15μM was proper concentrations for cell culture and tooth germ culture, respectively (S2 Fig A-C). The isolated E16.5 tooth germs were cultured in air-liquid interface culture system for 7 days, which made them develop into approximately E19.5 or P0 stage *in vitro*, and we found that tooth germ development was delayed by SP600125 treatment, especially the formation of enamel organ and dental papilla, under microscope. For example, the shape of enamel organ treated with SP00125 was more obscure than one without SP600125 (Fig 3, A-B, D-E). H&E staining of sample with DMSO showed the development stage of tooth germ as late bell stage, with plenty of polarized odontoblasts along basement membrane. But sample with 15μM SP600125 showed the development stage of tooth germ as early bell stage, with undifferentiated mesenchymal cells along BM (Fig 3, C, F).

**Fig 3.**
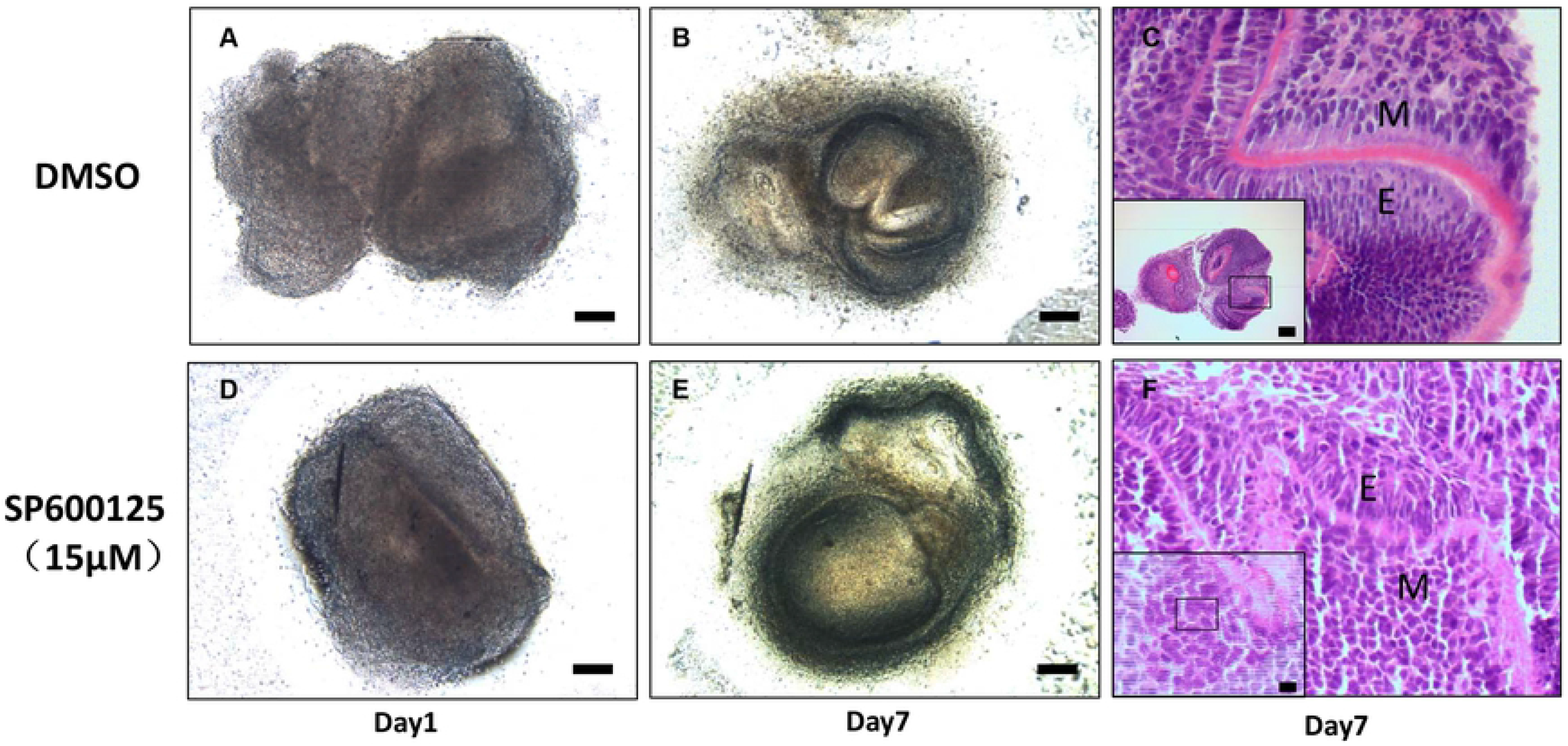
SP600125 postponed tooth germ development *in vitro*. (A, D) Organ culture with DMSO or SP600125(15μM) for 1 day, whole view of E16.5 tooth germs. (B,E) Organ culture with DMSO or SP600125(15μM) for 7 days, whole view of E16.5 tooth germs. (C) H&E staining of sample with DMSO showed the development stage of tooth germ as late bell stage, with plenty of polarized odontoblasts along basement membrane. (F) H&E staining of sample with 15μM SP600125 showed the development stage of tooth germ as early bell stage, with undifferentiated mesenchymal cells along basement membrane. Bars=200μm. M: mesenchyme, E: epithelium

### SP600125 inhibit cell migratory polarity and functional polarity by migration and differentiation assay

As tooth development and cell polarity formation were retarded with SP600125 treatment, we wondered if the migration and differentiation ability of dental papilla cells with SP600125 treatment was changed. With SP600125 treatment, the migration of mDPCs and A11 cells was reduced, and the number of cells at the leading edge with the Golgi polarized within a 120°arc in front of the nucleus were decreased (Fig 4, A, B). With SP600125 treatment, ALPase activity was reduced significantly in OM-induced mDPCs at day 5 and A11 cells at day 3. Mineralized nodules were decreased at day 14 and 7 respectively (Fig 4, C, D). These results suggested JNK signaling play roles on cell migration and differentiation, and SP600125 treatment inhibit the migratory polarity and functional polarity of mDPCs or A11 cells.

**Fig 4.**
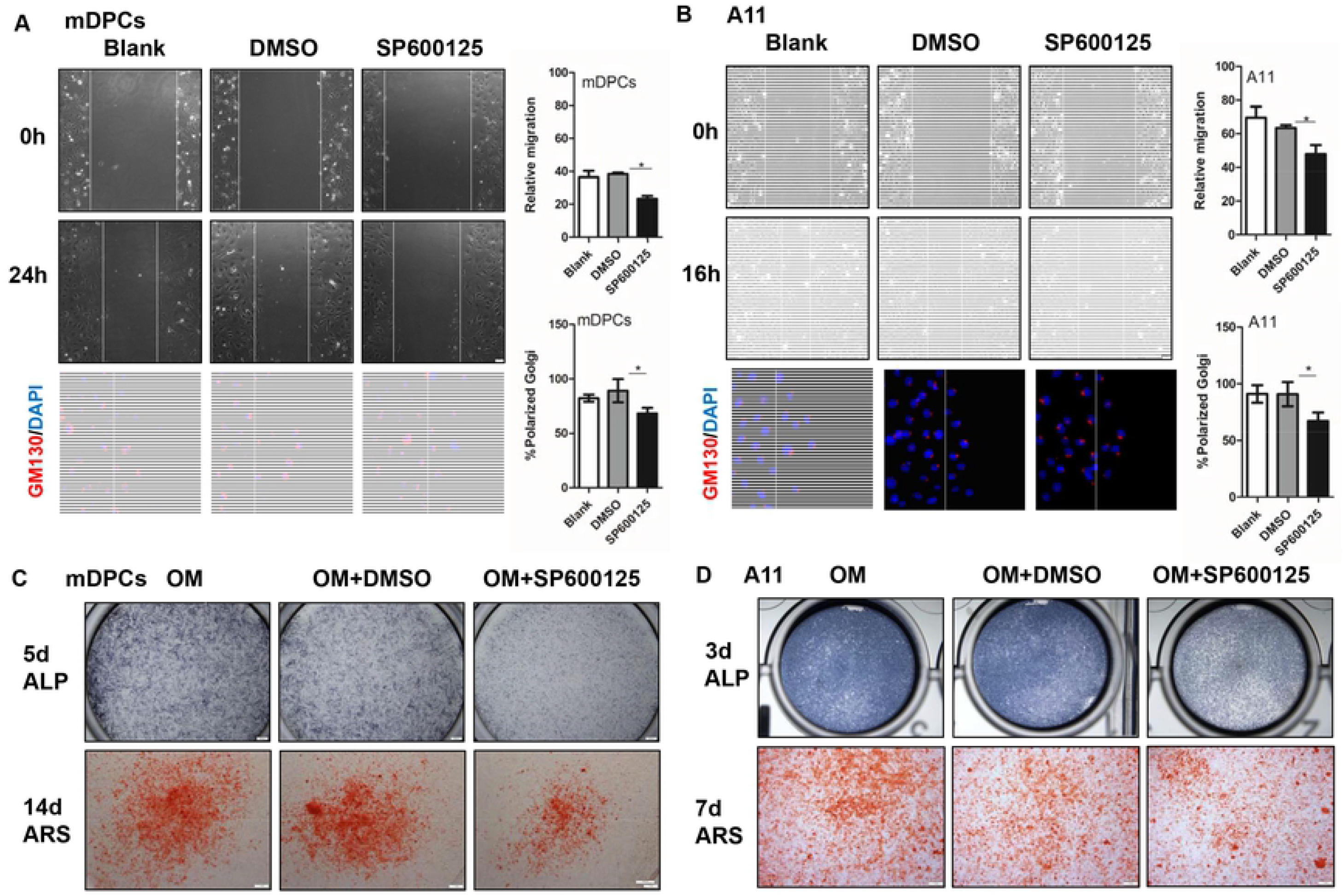
SP600125 reduced cell migration and differentiation. Scratch assay and cell immunofluorescence of GM130 in mDPCs(A) or A11 cells(B), results showed that 10μM SP600125 could inhibit cell migration and polarity formation along the scratch line. ALP staining and alizarin red staining in mDPCs(C) or A11 cells (D) in OM with 10μM SP600125 treatment, results showed that SP600125 treatment could inhibit cell differentiation. Bars: 50μm (A,B), 200μm (C,D) \**P*<0.05 with significance.

### Changes of some polarity-related genes in dental papilla tissues and cells

By gene screening, we found some polarity-related genes as Map1b, Rac1, Cdc42, RhoA, Celsr1, Snai1, Zeb2, Prickle3, Golga2 and Golga5 were up-regulated in dental papilla tissues of P1, compared with E17.5 (Fig 5, A). Meanwhile, the expression of these genes was detected in OM-induced mDPCs and A11 cells (Fig 5, B, C). Finally, we found Prickle3, Golga2, Golga5 and RhoA were up-regulated during the process of dental papilla development, as well as mDPCs and A11 differentiation (Fig 5, D). Furthermore, their expression was reduced accordingly in mDPCs transfected with SAPK/JNK siRNA or treated with SP600125(Fig 5, E). These results might remind possible role of Prickle3, Golga2, Golga5 and RhoA participating in JNK signal regulating pathway.

**Fig 5.**
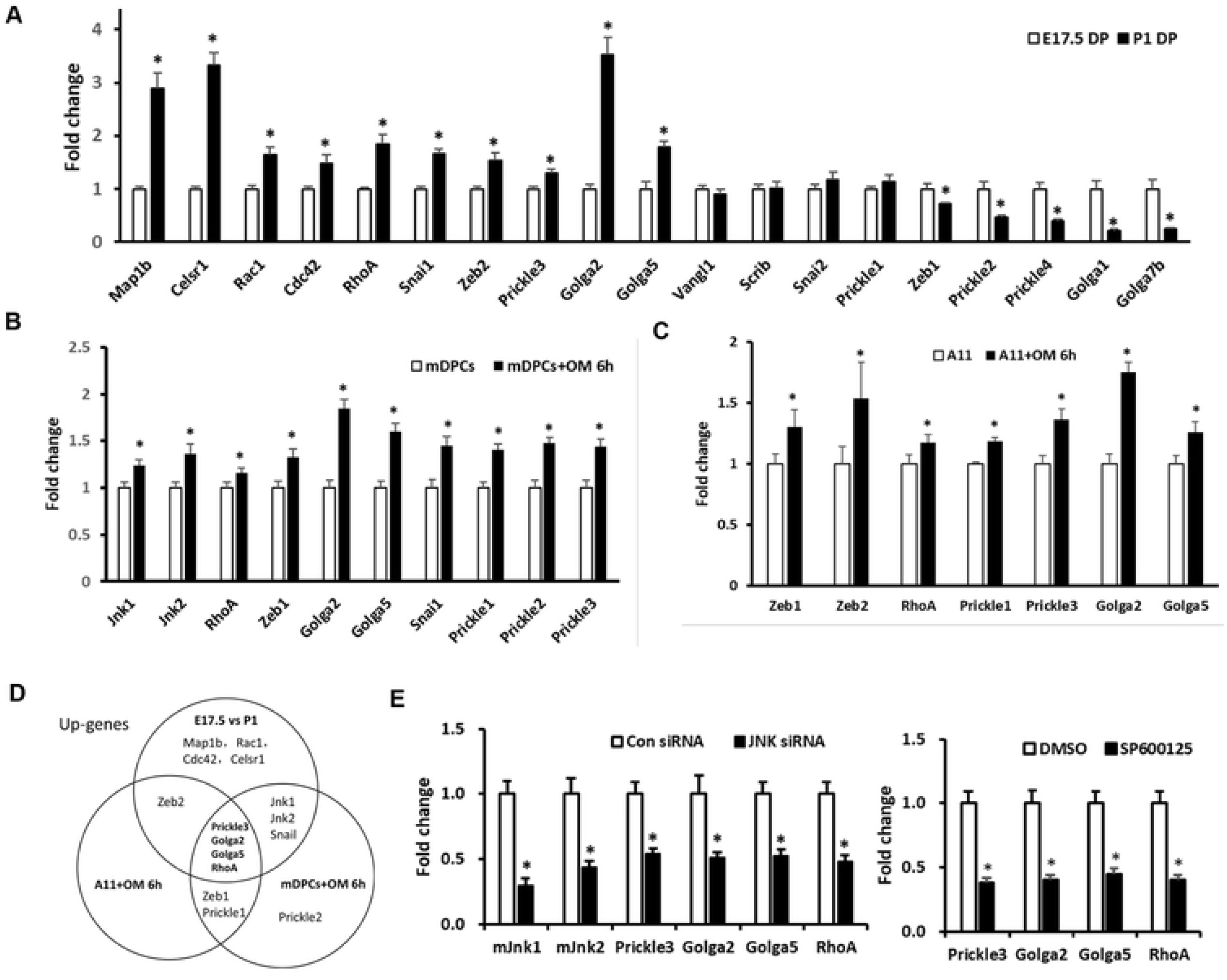
Polarity-related gene expression in dental papilla tissues and cells. (A) Expression of polarity-related genes in P1 dental papilla tissues compared with E17.5. (B) Up-regulated screened genes in mDPCs after osteogenic induction for 6h. (C) Up-regulated screened genes in A11 cells line after osteogenic induction for 6h. (D) Up-regulated genes both in A,B and C. (E) Expression of Prickle3, Golga2, Golga5 and RhoA in mDPCs with SAPK/JNK siRNA transfection or SP600125(10μM) treatment, results showed that similar changes between JNK siRNA and SP600125 treatment. DP: dental papilla, OM: Osteogenic medium, mDPCs: mouse dental papilla cells. \**P*<0.05 with significance

### RhoA signaling contributed to the JNK-regulated cell differentiation and polarity formation

As we sorted out some up-regulated polarity-related genes as Prickles, Golga2, Golga5 and RhoA, we chose RhoA gene and used Ad-RhoA WT and Ad-RhoA Q63L to transfect mDPCs in order to study the role of RhoA activity during this process, because our previous study found RhoA-JNK signaling pathway could affect human dental papilla cells adhesion, migration and differentiation[15], and we hypothesis that RhoA would have a role on JNK-regulated cell polarization. SP600125 could down-regulate RhoA activity significantly (Fig 6, A, lane 1 vs lane 2, lane 3 vs lane 4), and RhoA Q63L could up-regulate RhoA activity (Fig 6, A, lane 1 vs lane 3, lane 2 vs lane 4). Compare to RhoA WT, RhoA Q63L could significant up-regulate SP600125-inhibited RhoA activity (Fig 6, A), and partly rescue SP600125-inhibited mDPCs differentiation (Fig 6, B) and SP600125-inhibited mDPCs polarity formation (Fig 6, C). These data suggest that RhoA play a role on JNK signaling related cell polarization, which confirmed the results of polarity-related gene screening.

**Fig 6.**
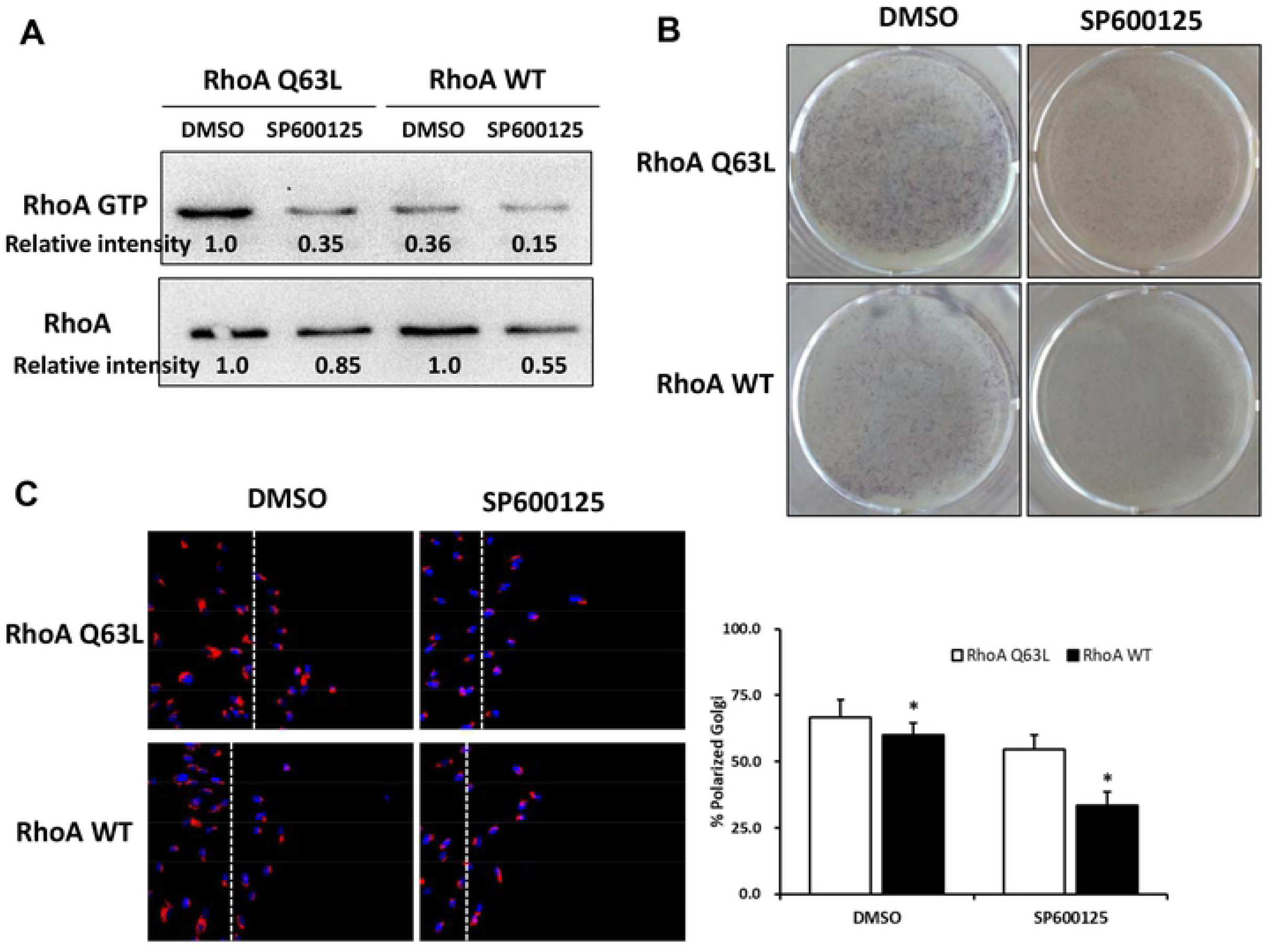
RhoA signaling contributed to the JNK-regulated cell differentiation and polarity formation. First, mDPCs were infected with RhoA mutant adenoviruses (RhoA WT or RhoA Q63L) for 48h, then were cultured with DMSO or SP600125(10μM). (A) After 24 h with SP600125 treatment, GST-Pull down assays were performed, RhoA Q63Lcould partly up-regulated the RhoA activity with SP600125 treatment. (B) After 5 days with SP600125 treatment, ALPase staining showed that RhoA Q63L could partly rescue SP600125-regulated inhibition of mDPCs differentiation. (C) After 10 h with SP600125 treatment, cell immunofluorescence of GM130 for the cells along scratch line were analyzed, and RhoA Q63L could partly rescue SP600125-regulated decreased of mDPCs polarity formation. \**P*<0.05 with significance

## Discussion

During the process of murine tooth development, the invaginating epithelium is surrounded by condensing mesenchyme transforming into bud stage (E12), following cap stage (E14), bell stage (E15), and morpho-differentiation (E17), dentinogenesis (P2)[24]. And mesenchymal cells facing the basement membrane will differentiate into dentin producing odontoblasts[25]. In this study, we choose several time points during morpho-differentiation stage, E17.5 represent the stage ready to cell differentiation, and P1 represent the stage close to finished cell differentiation. And our data also show similar tissue structure as describe[24], by HE staining, there are no significant cell morphology change and polarity formation in E17.5 tooth germ, but very significant cell polarity in P1 tooth germ. Tooth germ organ culture method was proven technique[26], and we had modified it to culture tooth germ and reconstituted tooth germ[19, 20]. By the way, if we try to study the role of some factors on tooth germ cell differentiation, we should give the treatment before morpho-differentiation stage, but as close to this stage as possible in order to avoid organ culture *in vitro* for too long time, so we chose E16.5 tooth germ to given extraneous treatment. Our study found that E16.5 tooth germ was develop to P0[27] or E19.5 stage tooth germ after 7 days organ culture *in vitro*. Above of all, E17.5 and P1 time points could be used to represent before and after dental papilla cells differentiated, and E16.5 tooth germ was suitable to study the change of dental papilla tissue changes by organ culture.

As early as 1993, polarizing odontoblasts have been described[1], dental papilla cell polarization process included cell polarization in migration process towards basement membrane (BM), and cell functional polarity with asymmetry in morphology to secret matrix. In cell migration process, polarity is conventionally referred to as asymmetric signaling activities and cellular structures along the front-rear axis. It is usually hard to define a solitary front-rear axis in a migrating cell, but easy to define an apical-basal axis[28], apical-basal axis usually exists in epithelial cells and functional cells like odontoblast, ameloblast, etc. In this study, we found that combining of scratch assay and GM130 staining was available to study the migratory polarization, and JNK inhibitor SP600125 treatment could reduce the migratory polarization of mDPCs.

About the role of JNK signaling on cell polarization, previous studies demonstrated the role of JNK signaling on cell polarity and mesenchymal fate[14, 29], and we also found JNK signaling had a role in odontoblasts maturation[15]. Some authors[30] have studied that role of JNK signaling in tissue morphology, and found that in cultured neurons from the intermediate zone, activated JNK was detected along microtubules in the process, and application of SP600125 caused irregular morphology and increased stable microtubules in neuron processes. It is a possibility of the involvement of JNK in controlling tubulin dynamics in migrating neurons[30]. In our study, combined with IHC, RT-PCR and WB results, all of them demonstrated JNK signaling was activated in the process of polarity formation of dental papilla cells, and SP600125 could induce some delayed effects, which inspired us to further explore the polarity-related mechanism.

Usually, cell polarity is defined as asymmetry in cell shape, protein distribution and cell function[31]. One characteristic of cell polarity is polarized distribution of cytoplasmic organelles. As a marker of cell polarity, Golgi complex function in a variety of membrane-membrane and membrane-cytoskeleton tethering events. GM130 (Golgi matrix protein of 130kDa), a static structural matrix associated with the Golgi apparatus, is important for protein transport, cell migration and polarization[32, 33]. So we chose GM130 as a maker to observe polarity change as previous studies[21, 22]. In tooth germ, IF data showed GM130 were laterally distributed in the functional side of odontoblasts, suggesting cell functional polarity. In cultured cells *in vitro*, cell polarity change was observed by quantitation of cells at the leading edge with Golgi polarized within a 120° arc in front of the nucleus as previously described[15, 21, 22], suggesting cell migratory polarity. Additionally, we examined the gene expression of Golga1, Golga2, Golga5 and Golga7b, the results showed the expression of Golga2, Golga5 were increased in P1 dental papilla compare to E17.5, but not Golga1 and Golga7b. As Golga2/GM130 mRNA and protein were higher expressed in P1 tooth germ, compared to E17.5, and literatures show Golga5, also known as Golgin 84, is transmembrane golgin and combined with Rab1, playing a key role in formation and maintenance of golgi apparatus[34]. In our study (figure 4 and 5), JNK inhibitor could disturb GM130 polar distribution, and SAPK/JNK siRNA infection could decrease the expression of Golga2 and Golga5 mRNA too. So, we speculate that JNK signaling plays a role in golgin expression and distribution, further effects cell polarity formation.

Another characteristic of cell polarity is polarity proteins. As characteristic polarity protein in odontoblasts is less known, we reviewed the literature and screened some polarity-related genes which are discussed as following. For example, Map1b and Celsr1 was increased in P1 dental papilla, which was verified by previous study about Map1b and Celsr1 distribution in odontoblasts[6, 8]. Prickle3, a planar cell polarity protein, was positive expression in papillary layer cells and some singular cells adjacent to the dentin[7]. RhoA, prototypical Rho GTPases, plays integral roles in cytoskeletal arrangement, membrane-trafficking pathways and ECM interactions which are crucial for cell polarization[35]. Interestingly, the expression of RhoA in our study was the same trend as Prickle3, and we had studied the role of RhoA on the migration of human dental papilla cells using RhoA mutant adenoviruses and found RhoA activity could active JNK signaling[15]. This can be easily explained as RhoA has function on cytoskeleton regulation and polarity formation.

RhoA, prototypical Rho GTPases, plays integral roles in cytoskeletal arrangement, membrane-trafficking pathways and ECM interactions which are crucial for cell polarization[35]. This can be easily explained as RhoA has function on cytoskeleton regulation and polarity formation. Hayes MN et al.[36] observed reductions of total and activated forms of RhoA in Vangl2-depleted rhabdomyosarcoma cells, and suggested RhoA functions downstream of Vangl2 to regulate cell growth and self-renewal. In our study, JNK inhibitor treatment also could down-regulate the expression of RhoA mRNA and inhibit RhoA activity. After SP600125 treatment, constitutively active RhoA (RhoA Q63L) could partly rescue SP600125-regulated inhibition of mDPCs polarity formation and differentiation. This result suggested that RhoA signaling contributed to the JNK-regulated cell differentiation and polarity formation of mDPCs. The role of Prickle3, Golga2, Golga5 in JNK-regulated cell polarity formation and differentiation still need further study.

Considering the accessibility and representation, in this study, we cultured poorly differentiated mDPCs, and highly differentiated odontoblast-like cell line A11 *in vitro*, which are represent different differentiation status of dental mesenchymal cells. The results from mDPCs and A11were consistent with that from dental papilla tissue, which enhanced the feasibility of using cultured cells *in vitro* for further research. During the process of dental papilla cell migrate towards BM, cell polarization should be formed, and our results of scratch assay showed that cell migration was inhibited by SP600125 treatment, and cell number at the leading edge with polarized Golgi was reduced. Also, during the process of dental papilla cell function and differentiation, cell polarization should be formed and cell morphology should be changed in tissue, so we tried to use the status of cell differentiation to reflect the degree of cell polarization formation *in vitro*. A11 cells were odontoblast-like cells, which can rapidly secret matrix vesicles after inducing by osteogenic medium[37]. SP600125 could inhibit ALPase activity and mineralized nodules formation in mDPCs and A11.Also, RhoA Q63L could partly rescue SP600125-regulated decreased of mDPCs polarity formation. So we illustrate SP600125 could influence polarity proteins and Golgi apparatus, restrain dental papilla cell polarity distribution and cell migration, which in turn affects cell differentiation even protein synthesis and secretion.

## Conclusion

JNK signaling has a positive role in dental papilla cells polarization. Prickle3, Golga2, Golga5 and RhoA are related with the mechanism of polarization, and RhoA signaling contributed to the JNK-regulated cell differentiation and polarity formation.

## Supporting information

**S1 Fig. Separated dental epithelium and papilla tissue.** (A, B) Separated dental epithelium and dental papilla under the stereomicroscope. (C) Expression of epithelium marker Cdh1 and mesenchymal marker Lhx8 in separated dental papilla. DP: dental papilla, DE: dental epithelium

**S2 Fig. Dose-response experiment of SP600125 on cell culture and tooth germ organ culture *in vitro*.** (A) For mDPCs culture, 10μM SP600125 could inhibit JNK signaling activation. (B) For tooth germ culture, 15μM SP600125 could inhibit JNK signaling activation. (C) For tooth germ cultue, 25μM SP600125 have some toxic effects on tissues, but not 15μM SP600125. Bars=200μm

**S1 Table. Primers sequence from Primerbank.**

